# Pan-cancer remodeling of reactive oxygen species homeostasis: distinct roles of the ROS-generation and antioxidant arms in immune context and outcome

**DOI:** 10.64898/2026.10.10.758205

**Authors:** Kun Xie, Yulong Xuan, Rongjun Sun, Ruyin Chang, Jiajing Lin, Youqi Shen, Wenyue Jin, Qi Lu, Haifeng Lu, Qunlin Zhang

## Abstract

Reactive oxygen species (ROS) are double-edged in cancer: tumor cells sustain high ROS generation while up-regulating antioxidant defense, so that the net balance between the two arms, rather than absolute ROS abundance, is the biologically meaningful quantity. Whether that balance defines a conserved tumor state across cancer types has been unclear. Here an experimentally measured ROS phenotype, obtained from a three-channel gold-nanocluster array, was generalized to public transcriptomes through a 107-gene ROS program scored in a large cell-line panel and in pan-cancer tumor and adjacent-normal tissue. The optical readout was reproducible and fully discriminative between cell lines, establishing the fingerprint as a quantitative starting point. In tumors, the generation and antioxidant arms were coordinately up-regulated relative to normal tissue, yet the two arms dissociated in their biology. The composite balance score tracked reduced immune infiltration consistently across cohorts, but arm-level decomposition showed that it was carried by low generation-arm activity in immune-cold tumors and followed arithmetically from how the balance score is defined; the association does not implicate antioxidant defense in immune infiltration. The clinical association pointed the other way: adverse outcome was attributable to the antioxidant arm, whereas generation was null. In cohort-held-out validation, tumors were separated from adjacent normal tissue, with redox programs outperforming generic phenotypic programs, indicating that the same program transfers across tumor types. Tumor remodeling of ROS homeostasis is therefore widespread, but its two arms carry distinct meanings: the balance score, driven by its generation term, reports immune context, whereas the antioxidant arm reports clinical risk.

## 1. Introduction

Cancer remains a leading cause of death worldwide ^[1]^, and reactive oxygen species (ROS) are central to its biology. Elevated ROS generation promotes DNA damage, genomic instability and oncogenic signaling, yet the same species can trigger oxidative cell death and senescence, which is the rationale for ROS-elevating therapies ^[2-5]^. Tumor cells resolve this paradox by simultaneously expanding their antioxidant capacity: NRF2-dependent programs and the cystine transporter SLC7A11/xCT sustain glutathione synthesis, and thioredoxin and peroxiredoxin systems buffer peroxides ^[6-10]^.

Because generation and scavenging are co-regulated, absolute ROS abundance is an unstable readout of tumor state. The net ROS balance - the relative strength of antioxidant defense versus ROS production - is more informative, and associations of this balance with epithelial-mesenchymal transition, stemness, ferroptosis susceptibility and immune evasion have been reported ^[7,9-14]^. However, most evidence comes from single cancer types, from individual genes, or from cell models, and a systematic pan-cancer description of the tumor ROS state is missing.

In previous work, pH-tuned histidine-templated gold nanoclusters (AuNCs@His) were developed that produce a three-channel fluorescent response to ROS, and the resulting response fingerprints were used to distinguish cancer cell types and proliferation states in vitro ^[15,16]^. That work established that ROS chemistry can be read out optically from living cells, but it was limited to a small number of cell lines and was not connected to tumor genomics or to clinical outcome.

One central question is addressed here: whether a measurable ROS phenotype reveals a conserved tumor state across cancer types. The optically read ROS fingerprint is first examined for quantitative robustness and for its capacity to discriminate tumor from non-tumor cells. Next, the extent to which ROS homeostasis is remodeled in a common direction across cancer types is assessed, together with the tumor programs that accompany such remodeling. Finally, the immunological and prognostic correlates of the tumor ROS state are examined, including the transferability of the underlying program across cancer types by classification. These analyses combined the array data with large public transcriptomic resources (CCLE, TCGA), using a 107-gene ROS program score and strictly cohort-separated validation (Figs 2-6 and Fig. S1).

**Figure 1.**
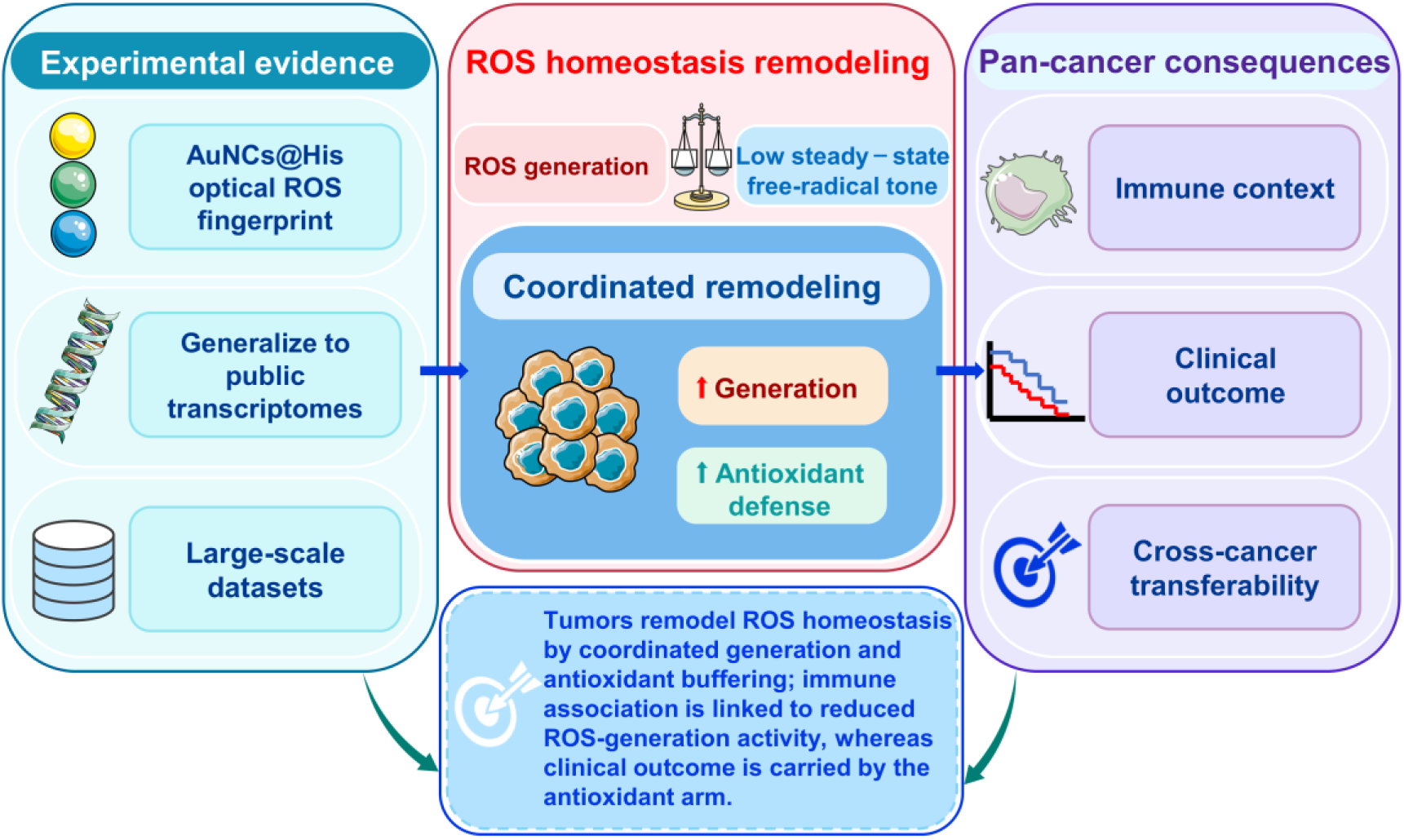
Overview of the study design and main findings. Concept hierarchy linking the optical ROS fingerprint to tumor biology. Left, experimental evidence: the AuNCs@His three-chan nel array provides an experimentally measurable ROS phenotype (11 cell lines, six ROS species), which is generalized to public transcriptomes through a 107-gene ROS program (generation 33, antioxidant defense 39, proliferation 15, EMT 10, stemness 10) and scored across 33 TC GA cohorts (9,732 tumors and 727 adjacent normals). Center, ROS homeostasis is remodeled - coordinated increases in generation and antioxidant defense - rather than uniformly elevated. Right, pan-cancer consequences: immune context, clinical outcome and cross-cancer transferability. Graphical elements in this figure were adapted from Servier Medical Art (https://smart.servier.com), licensed under CC BY 4.0 (https://creativecommons.org/licenses/by/4.0/).

**Figure 2.**
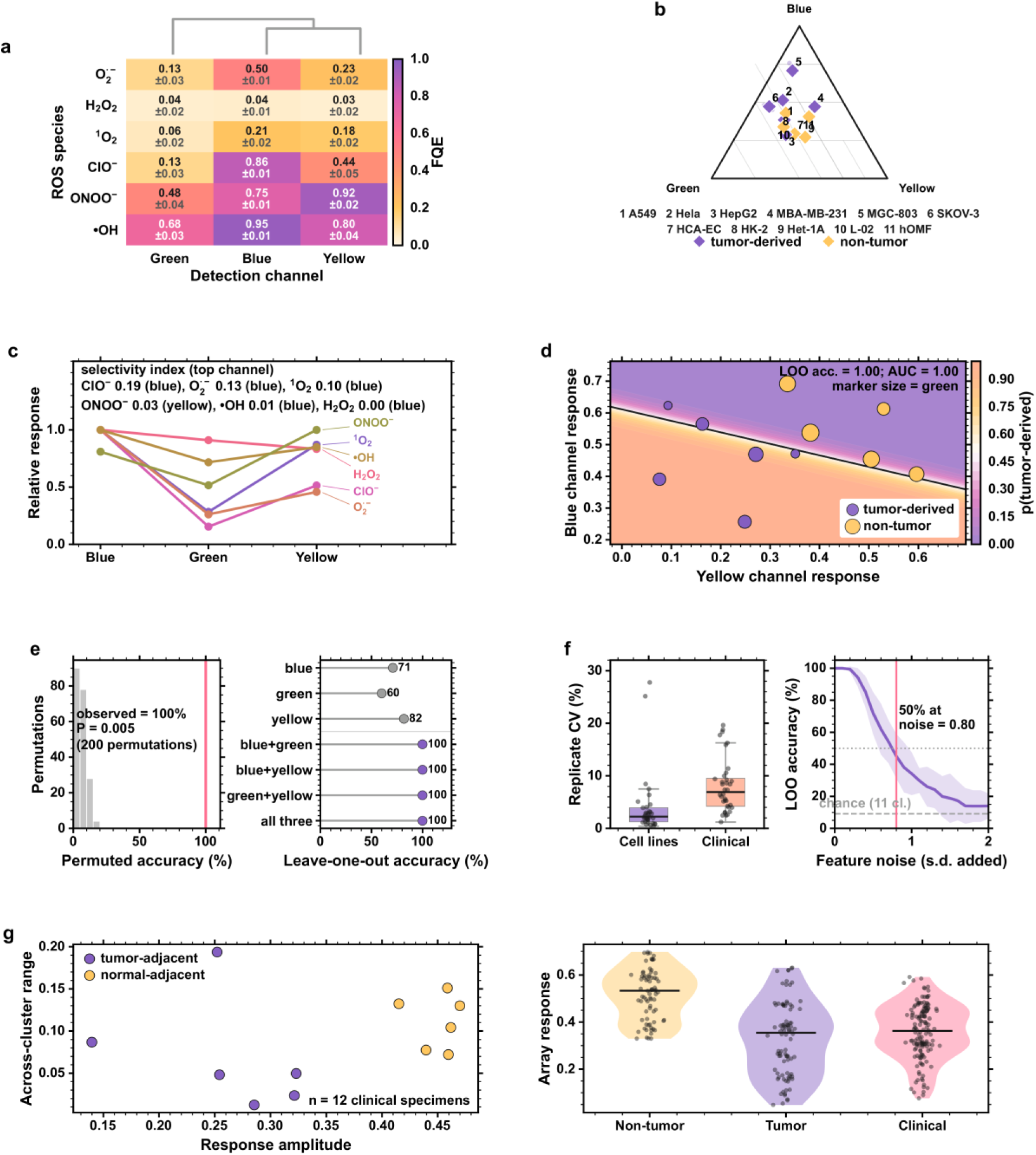
Optical ROS fingerprinting provides a reproducible phenotypic readout. (a) Response matrix of the three AuNCs@His channels (blue, green, yellow) to six ROS species (ClO^−^, ONOO^−^, ^•^OH, H_2_O_2_, ^1^O_2_, O_2_^•−^), shown with the s.d. of five replicates; rows and columns are ordered by average-linkage hierarchical clustering and the corresponding dendrograms are shown. (b) Fingerprint geometry of the 11 cell lines displayed as a ternary plot in the blue-green-yellow composition space: each line is shown as a diamond labeled with its key number (key below the panel), the pale points are the five technical replicates and the faint gray grid marks 25%, 50% and 75% composition. Tumor-derived lines (1-6: A549, HeLa, HepG2, MBA-MB-231, MGC-803, SKOV-3) show a higher blue fraction on average (0.47 versus 0.35) and non-tumor lines (7-11: HCA-EC, HK-2, Het-1A, L-02, hOMF) a higher yellow fraction (0.31 versus 0.20), with overlap between the groups; the five replicates of each line coincide, and the two groups are fully separated by their raw channel responses in the blue-yellow plane of panel d. (c) Channel-preference profiles of the six ROS species (responses normalized to each species’ maximal channel response), annotated with the selectivity index (1 - normalized entropy) and the preferred channel: ONOO^−^ and ClO^−^ are preferentially reported by the yellow and blue channels, respectively, whereas H_2_O_2_ has the flattest profile. (d) Leave-one-out discrimination of tumor-derived versus non-tumor cell lines, shown as a decision map in the blue-yellow channel plane: the background is the LDA posterior probability (color scale), the black line is the 0.5 boundary, marker size scales with the green channel response, and accuracy is 1.00 (n = 55 samples). (e) Permutation null distribution of the 11-class classifier (200 label permutations; observed accuracy 100%, P = 0.005) and leave-one-out accuracy for channel subsets: the three single channels (blue, green, yellow; 60-82%), the three channel pairs and all three channels (100%), showing the intrinsic redundancy of the three-channel array. (f) Robustness of the readout: coefficient of variation of technical replicates for the cell-line and clinical datasets, and degradation of the leave-one-out accuracy under multiplicative feature noise (mean with 95% interval over repeated noise draws; the dashed line is chance level for 11 classes); accuracy falls to 50% only at a noise level of about 0.80 s.d. (g) Quantitative description of the 12 clinical specimens (invasive breast cancer samples originally reported in [15]): response amplitude versus across-cluster range, and comparison of the non-tumor and tumor cell-line responses with the clinical specimens (box, median).

## 2. Materials and methods

### 2.1 Cellular ROS-response data

ROS-response fingerprints were generated with the three-channel AuNCs@His array described previously [15]; both the cell-line and the clinical response data analyzed here are the measurements originally published in that study. Briefly, three pH-tuned nanocluster preparations (response clusters denoted by color: blue, green and yellow) were incubated with six ROS species (ClO^−^, ONOO^−^, ^•^OH, H_2_O_2_, ^1^O_2_ and O_2_^•−^), and fluorescence quenching efficiencies (FQE) were computed as FQE = (F_0_ - F_1_) / F_0_, where F_0_ and F_1_ are the emission intensities before and after ROS addition. The resulting response matrix comprises 11 cell lines (A549, HCA-EC, HK-2, HeLa, HepG2 and six additional lines; five biological replicates each, 55 samples in total) and 12 clinical specimens, the latter being the 12 cases of invasive breast cancer samples described in that study, collected from the First Affiliated Hospital of Anhui Medical University (the ethics approval under which they were obtained is given in the Declarations). Cell lines were grouped into tumor-derived and non-tumor groups according to their origin.

### 2.2 Public transcriptomic data

Pan-cancer tumor and adjacent-normal expression data were obtained from the UCSC Xena browser as TCGA HiSeqV2 log2(normalized count + 1) matrices for 33 cohorts (10,459 samples: 9,732 tumors and 727 adjacent normals) ^[17,18]^. Sample type was assigned from the TCGA barcode; the primary analysis used primary-tumor samples only (sample code 01) and adjacent normals (code 11), and samples with other tumor codes (02, 03, 05, 06, 07; 663 samples in the full phenotype table, 5.2%) were excluded, as were metastatic and recurrent specimens. Clinical and survival annotation (overall survival, OS; progression-free interval, PFI; disease-specific survival, DSS) was taken from the Xena pan-cancer survival table. Cell-line expression data were obtained from the Cancer Cell Line Encyclopedia (CCLE RNA-seq, RSEM TPM; 1,019 cell lines annotated to 14 lineages) ^[19,20]^. All public datasets were used without re-processing beyond the transformations described below.

### 2.3 ROS program gene sets and scoring

A 107-gene ROS program comprising five sets was defined: ROS generation (33 genes), antioxidant defense (39), proliferation (15), epithelial-mesenchymal transition (EMT, 10) and stemness (10) (Table S2 lists the ROS-program genes with their program membership; Table S5 also lists the immune program and the 13 checkpoint and effector genes). Two of the 107 entries are alternative symbols — NOX2 for CYBB and PIG3 for TP53I3 — so the program corresponds to 105 unique genes; the alias entries are absent from the expression matrices and contribute neither to program scores nor to the classifier features. Genes were taken from canonical, literature-curated catalogs of ROS-producing enzymes (NADPH oxidases and their regulatory subunits, mitochondrial and peroxisomal sources, oxidases), antioxidant enzymes and NRF2-dependent genes, and from established proliferation, EMT and stemness marker sets; the sets were fixed before scoring and were not optimized on any outcome. All genes were weighted equally, and sensitivity to this choice was examined with ranking-based (median z-score) and leave-one-gene-out scoring. Partial collinearity with proliferation and EMT is expected by construction, since the program deliberately includes those axes; the ablation analysis in Section 3.5 quantifies what each program contributes individually, and the random-set comparison provides the corresponding null. For every dataset, expression values were z-scored per gene within cohort (so that cohort-specific and platform-specific offsets are removed), and a program score was the mean z-score of its member genes. Because z-scoring uses the distribution of the cohort being scored, scores in tumor-versus-normal tasks are not purely held-out quantities, and absolute values are not comparable between cohorts; cross-cohort inferences here rest on within-cohort effect directions and on meta-analysis, not on pooled absolute values. The ROS balance score was defined as antioxidant score minus generation score, so that positive values denote an antioxidant-shifted state. An immune program score (20 interferon/cytotoxic genes) and 13 individual immune-checkpoint or immune-effector genes (PDCD1, CD274, CTLA4, LAG3, HAVCR2, TIGIT, IDO1, CD8A, IFNG, GZMA, CXCL9, B2M, HLA-A) were analyzed with the same scoring rules.

### 2.4 Array fingerprint analyses

Response matrices were clustered hierarchically (average linkage on correlation distance). ROS selectivity was quantified per species as 1 - normalized Shannon entropy of its response across the three channels, together with the ratio of the strongest to the second-strongest channel response. Discriminative performance was assessed by leave-one-out cross-validation of four classifiers (LDA, SVM with RBF kernel, kNN with k = 3, random forest), by permutation testing (200 label permutations), by repeated cross-validation of technical replicates, by channel ablation and by noise titration of the feature matrix. Binary discrimination of tumor-derived versus non-tumor cells was evaluated by leave-one-out cross-validation in the two-dimensional blue-yellow plane, and the classification result was visualized as a decision map together with the area under the ROC curve.

### 2.5 Cell-line analyses

Program scores were computed for 1,019 CCLE cell lines. Lineage-specific deviations of the balance score were tested against all other lineages (two-sided Mann-hitney U test with Benjamini-Hochberg correction). Pairwise associations between the balance score and the proliferation, EMT and stemness programs were quantified with Pearson and Spearman correlation; effect sizes are reported with Fisher-z 95% confidence intervals.

### 2.6 Pan-cancer analyses

For each TCGA cohort, differences in program scores between tumors and adjacent normals were tested with two-sided t-tests (Welch) and corrected with the Benjamini-Hochberg procedure across cohorts. Tumor-only pooled correlations between programs were computed across all 9,732 tumors, and per-cohort correlations were summarized as pooled coefficients, medians and the number of cohorts reaching FDR < 0.05.

### 2.7 Survival analyses

Survival analyses used tumor samples only (n = 9,559 with OS annotation). The balance score was dichotomized at the pan-cancer median for Kaplan-Meier estimates and log-rank tests, and cohort-wise univariate Cox models were fitted with the continuous per-s.d. score. Cohort-level hazard ratios were combined by fixed-effect inverse-variance meta-analysis for OS, PFI and DSS. Sensitivity to dichotomization was assessed with median splits, tertile comparisons (T3 versus T1) and continuous models. Analyses stratified by the median of the immune program score were performed to test whether the prognostic association depended on immune context.

### 2.8 Classifier development and validation (transferability test)

Tumor-versus-normal classification tested whether the ROS program carries a state that transfers across cancer types. Classification used the 107 ROS program genes as features, z-scored per gene within cohort, in the 24 cohorts containing at least 20 annotated samples of both classes (8,941 samples: 8,214 tumors and 727 adjacent normals). Five models (logistic regression with L2 penalty, L1/LASSO logistic regression ^[21]^, SVM with RBF kernel, random forest and gradient boosting ^[22]^) were compared by stratified 5-fold cross-validation repeated three times. External validation used leave-one-cohort-out training, in which the model was fitted on 23 cohorts and evaluated on the held-out cohort; predictions were pooled for overall ROC analysis and summarized per cohort. A fixed feature set chosen using all cohorts would leak information into the held-out evaluation. External validation used the full pre-specified 107-gene program, with no data-driven feature selection. The LASSO selection described below served only as a stability analysis of the program itself and was never model input for any reported AUC. All feature-selection and model-fitting steps were confined to the training cohorts; per-gene standardization was performed within each cohort before splitting and therefore also uses the distribution of the held-out cohort, so these predictions are cohort-held-out rather than strictly held-out quantities (Section 2.3). Feature stability was assessed by LASSO logistic regression over 100 bootstrap resamples (80% of samples). For reference, 100 random gene sets of the same size (107 genes) were drawn from 2,000 randomly selected background genes and evaluated with the same cross-validation scheme. Discrimination was summarized by AUC, and calibration by reliability curves and the Brier score (with Murphy decomposition) ^[23]^. Clinical utility was assessed by decision curve analysis ^[24]^.

### 2.9 Antioxidant sub-module and arm decomposition

To separate the contributions of the two arms of ROS homeostasis, the 39 antioxidant genes were partitioned into nine non-overlapping sub-modules defined by biochemical function (GSH synthesis and recycling; GSH peroxidases; catalase and SOD; thioredoxin-peroxiredoxin system; NRF2-KEAP1 axis; cystine transport (SLC7A11/SLC3A2); NADPH regeneration; iron storage; aldo-keto reductases; Table S3). Sub-module scores were the mean z-score of their members within cohort and were correlated (Pearson) with the immune program score in each cohort; direction consistency across cohorts was assessed with a two-sided sign test with Benjamini-Hochberg correction. Arm-level prognostic associations were quantified with univariate Cox models per standard deviation for the generation score, the antioxidant score and the composite balance score in tumors with survival annotation (Fig. S1g,h; Table S4).

### 2.10 Statistics

Unless stated otherwise, tests were two-sided and P values were corrected with the Benjamini-Hochberg false-discovery-rate procedure within each analysis ^[25]^. Effect sizes are reported with 95% confidence intervals. Kaplan-Meier estimates and log-rank tests followed standard definitions ^[26,27]^. Results that did not satisfy pre-specified strength criteria (statistical significance after correction, interpretable effect size, robustness to parameter changes and reproducibility across cohorts) were documented internally and excluded from this manuscript.

### 2.11 Software

Analyses were performed in Python 3.14 with NumPy, SciPy ^[28]^, pandas, scikit-learn^[29]^ and statsmodels; figures were produced with Matplotlib ^[30]^ and seaborn ^[31]^ under a single project-wide style configuration. All analysis scripts, intermediate data tables and figure-generating code are available from the corresponding authors on reasonable request.

## 3. Results

### 3.1 Optical ROS fingerprinting provides a reproducible phenotypic readout

The extent to which ROS chemistry can be read out from cells in a quantitative and reproducible manner was examined first, using the response matrix of the AuNCs@His array (11 cell lines and six ROS species, five replicates; Fig. 2a). Hierarchical clustering of response profiles separated the six ROS species into three response groups with systematic differences in channel preference (Fig. 2a). In the channel composition space, the individual cell lines occupied characteristic positions: tumor-derived lines had a higher blue fraction on average (0.47 versus 0.35) and non-tumor lines a higher yellow fraction (0.31 versus 0.20), with substantial overlap between the two groups, while the five technical replicates of each line coincided (Fig. 2b). Selectivity across the three channels was highest for ClO^−^ (blue channel; selectivity index 0.19) and O_2_^•−^ (0.13) and lowest for H2O2 (< 0.01), which gave an almost flat response profile; ONOO^−^ was set apart not by selectivity (0.03) but by its preferred channel (yellow) (Fig. 2c).

Discriminative performance was evaluated by leave-one-out cross-validation. All 55 samples were assigned to their 11 cell lines by all four classifiers (accuracy 100%, 95% exact binomial CI 93.5-100%; macro-F1 = 1.00), well above the permutation null (mean accuracy 0.057; 95th percentile 0.127; 200 permutations; P = 0.005; Fig. 2e). With five replicates and six species per line, however, this comparison rests on 11 distinct profiles and reflects separability of the panel rather than general classification accuracy; the full confusion matrix and per-model metrics are given in Table S6. After collapsing of the cell lines into tumor-derived and non-tumor groups, the two groups were separated without overlap in the blue-yellow channel plane (accuracy 1.00; Fig. 2d), and the decision surface remained well separated from the data points. Channel ablation indicated intrinsic redundancy: accuracies of 60-82% were retained with single channels, whereas complete discrimination was restored by any two channels (Fig. 2e). Technical reproducibility was high (median coefficient of variation 2.2% for cell-line replicates and 6.9% for clinical specimens), and the accuracy of the discrimination decreased to the 50% level only when the features were perturbed with about 0.80 s.d. of multiplicative noise (Fig. 2f). Response amplitudes and across-cluster ranges varied among the 12 clinical specimens, which were placed between the tumor and non-tumor cell-line distributions (Fig. 2g).

These observations indicate that the optical ROS fingerprint is a stable, low-dimensional and highly discriminative readout of cellular ROS chemistry, and provide a quantitative starting point for examining tumor ROS states across cancer types.

### 3.2 ROS homeostasis captures tumor-cell phenotypic heterogeneity across 1,019 cancer cell lines

For connection of the array readout to molecular programs at scale, the 107-gene ROS program was scored in CCLE cell lines (Fig. 3). In cell lines, the generation and antioxidant scores were only weakly coupled (r = 0.11, P = 6.6e-04; Fig. 3a), in contrast to the stronger coupling found in tumors (Section 3.3). The balance score differed by lineage, with significant deviation from the remaining lineages in 8 of 14 lineages after FDR correction (Fig. 3b), the largest shifts being found in hematopoietic and central-nervous-system lineages.

**Figure 3.**
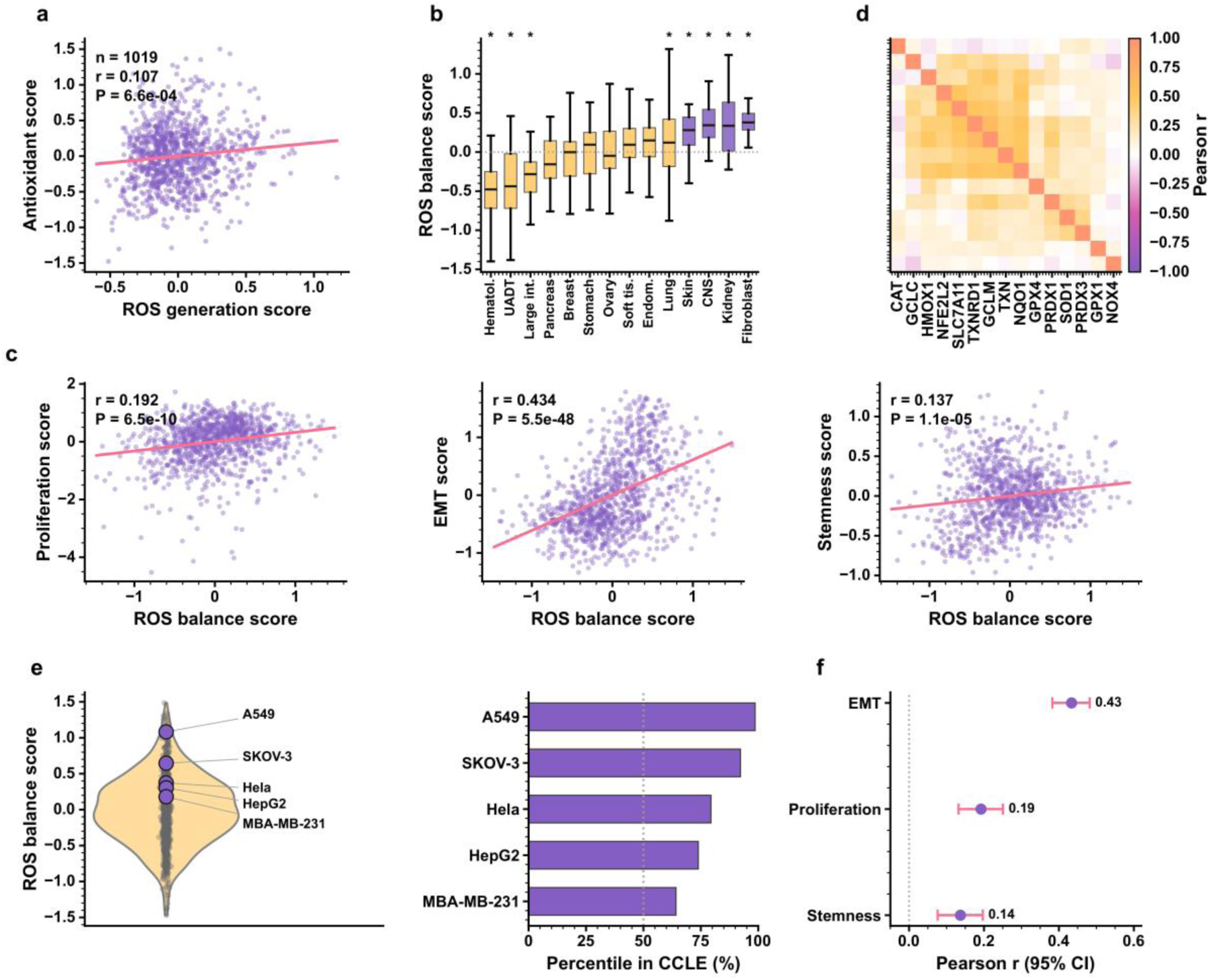
ROS homeostasis captures tumor-cell phenotypic heterogeneity across 1,019 cancer cell lines. (a) ROS generation versus antioxidant program scores in CCLE cell lines (Pearson r = 0.11, P = 6.6e-04). (b) Lineage distribution of the ROS balance score; lineages significantly different from the remaining lineages (Mann-Whitney U with Benjamini-Hochberg correction, FDR < 0.05) are highlighted (8 of 14 lineages). (c) Association of the balance score with proliferation, EMT and stemness programs (r = 0.19, 0.43 and 0.14, respectively; n = 1,019). (d) Co-expression structure of key redox genes across cell lines. (e) Descriptive placement of the cell lines used in the array experiments within the CCLE ROS-balance distribution (percentile ranks; only lines with a public counterpart are shown and no significance is claimed). (f) Summary of effect sizes (Pearson r with 95% confidence intervals, Fisher-z transformation).

Positive associations of the balance score were found with the EMT (r = 0.43, P = 5.5e-48, n = 1,019), proliferation (r = 0.19, P = 6.5e-10) and stemness (r = 0.14, P = 1.1e-05) programs (Fig. 3c,f), and the expected internal structure of the antioxidant arm (SOD1/CAT/GPX/PRDX families) was preserved across lineages in co-expression analysis (Fig. 3d). The cell lines used in the array experiments were placed within the CCLE distribution as a descriptive anchor (Fig. 3e); because a public counterpart was available for only a minority of these lines, this comparison is reported descriptively and is not used for inference.

In this independent cell-line panel, the ROS state co-varies with lineage and with programs related to tumor cell plasticity. As shown in Section 3.3, the direction of the EMT and stemness associations differs in intact tumor tissue, indicating context dependence of these phenotypic correlates.

### 3.3 Pan-cancer tumors undergo coordinated remodeling of ROS homeostasis

Scoring of the ROS program was next performed in 10,459 TCGA samples from 33 cancer types (9,732 tumors and 727 adjacent normals; Figs. 4 and S1a-c). Evaluable adjacent normal tissue (at least five adjacent normals) was available for 18 cohorts, which were used for tumor-normal contrasts. Tumor-normal differences were tumor-type dependent but involved both arms of ROS metabolism: significant alterations were found for ROS generation in 11 of the 18 evaluable cohorts (FDR < 0.05), for antioxidant defense in 8, and for the composite balance in 13, of which 9 were shifted toward antioxidant defense and 4 toward ROS generation (Fig. 4a; Fig. S1a; Table 1 and Table S1). Tissue-dependent direction of the shift - antioxidant-shifted squamous, gastric and lung tumors alongside generation-shifted thyroid and renal tumors - indicates that remodeling, rather than uniform elevation, is the common feature (Fig. 4b). Five of these 18 cohorts had fewer than 20 adjacent normal samples (Table 1); for these, tumor-normal contrasts are reported for completeness but are not interpreted individually, and they contribute little weight to the pooled pattern.

**Figure 4.**
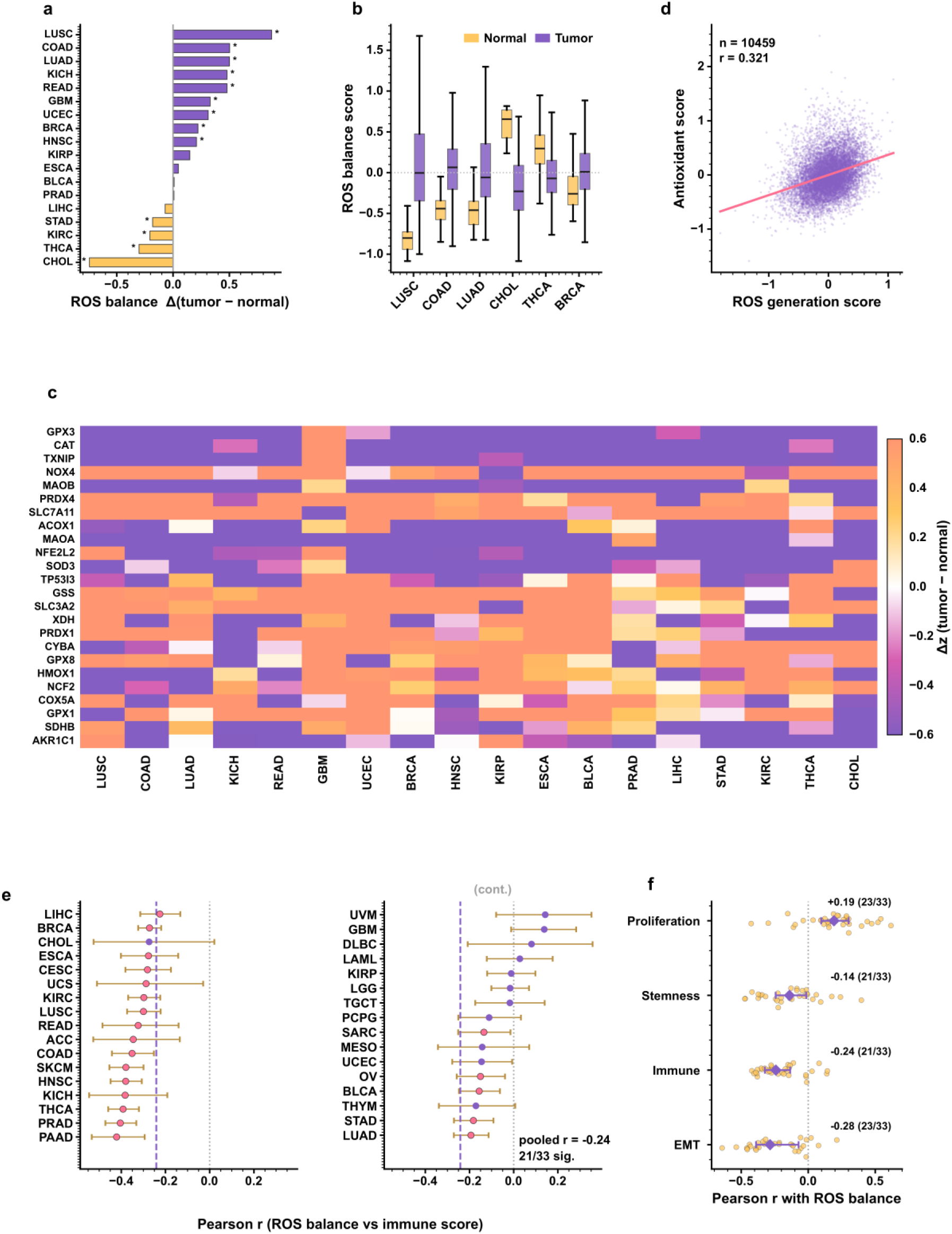
Pan-cancer tumors undergo coordinated remodeling of ROS homeostasis. (a) Tumor minus adjacent-normal difference in the ROS balance score per TCGA cohort (18 cohorts with evaluable adjacent normal tissue; n = 10,459 samples; cohorts ordered by effect size; filled markers denote FDR < 0.05; significant in 13 cohorts, of which 9 were antioxidant-shifted and 4 generation-shifted). (b) Representative distributions of the balance score in tumors and adjacent normals for selected cohorts. (c) Heat map of per-gene tumor-normal differences (delta z) across cohorts for the 24 most variable program genes (of the 70 measured in every cohort), showing concordant up- and down-regulation within each program. (d) ROS generation versus antioxidant program scores across 9,732 tumors (r = 0.33). (e) Per-cohort correlation between the balance score and the immune program score (pooled r = -0.24; FDR < 0.05 in 21 of 33 cohorts, direction negative in 29). (f) Pooled and per-cohort effect sizes of the balance score with proliferation, EMT, stemness and immune programs; the associations with EMT, stemness and proliferation are context dependent (compare Fig. 3c), whereas the inverse association with immune programs is consistent across cohorts.

**Table 1.** TCGA cohorts, sample numbers and tumor-normal difference in the ROS balance score (s.d.). Cohorts are ordered alphabetically; FDR values derive from two-sided Welch t-tests with Benjamini-Hochberg correction across cohorts; cohorts without evaluable adjacent normal tissue are marked n.a.

| Cancer type<br>(TCGA code) | Tumors (n) | Adjacent normals<br>(n) | ROS balance delta<br>(s.d.) | FDR |
| --- | --- | --- | --- | --- |
| ACC | 79 | 0 | n.a. | n.a. |
| BLCA | 407 | 19 | 0.0097 | 0.8912 |
| BRCA | 1104 | 114 | 0.2215 | 5.67e-13 |
| CESC | 305 | 3 | n.a. | n.a. |
| CHOL | 36 | 9 | -0.7425 | 0.0001 |
| COAD | 288 | 41 | 0.5026 | 3.99e-15 |
| DLBC | 48 | 0 | n.a. | n.a. |
| ESCA | 185 | 11 | 0.0474 | 0.8912 |
| GBM | 167 | 5 | 0.3297 | 0.0259 |
| HNSC | 522 | 44 | 0.2081 | 0.0038 |
| KICH | 66 | 25 | 0.4794 | 1.61e-06 |
| KIRC | 534 | 72 | -0.2058 | 6.25e-08 |
| KIRP | 291 | 32 | 0.1490 | 0.2135 |
| LAML | 173 | 0 | n.a. | n.a. |
| LGG | 530 | 0 | n.a. | n.a. |
| LIHC | 373 | 50 | -0.0702 | 0.3176 |
| LUAD | 517 | 59 | 0.5010 | 1.84e-17 |
| LUSC | 502 | 51 | 0.8742 | 1.70e-25 |
| MESO | 87 | 0 | n.a. | n.a. |
| OV | 308 | 0 | n.a. | n.a. |
| PAAD | 179 | 4 | n.a. | n.a. |
| PCPG | 184 | 3 | n.a. | n.a. |
| PRAD | 498 | 52 | 0.0083 | 0.8912 |
| READ | 95 | 10 | 0.4792 | 0.0002 |
| SARC | 263 | 2 | n.a. | n.a. |
| SKCM | 473 | 1 | n.a. | n.a. |
| STAD | 415 | 35 | -0.1813 | 0.0089 |
| TGCT | 156 | 0 | n.a. | n.a. |
| THCA | 513 | 59 | -0.3020 | 9.08e-11 |
| THYM | 120 | 2 | n.a. | n.a. |
| UCEC | 177 | 24 | 0.3120 | 7.20e-06 |
| UCS | 57 | 0 | n.a. | n.a. |
| UVM | 80 | 0 | n.a. | n.a. |

In tumors, the generation and antioxidant scores were strongly positively coupled (r = 0.33 across 9,732 tumors; Fig. 4d and Fig. S1c), three times the coupling seen in cell lines (r = 0.11), consistent with coordinated induction of both arms in the tumor context. Associations of the balance score with tumor-state programs were context dependent: in tumor tissue, inverse associations were found with EMT (r = -0.28) and stemness (r = - 0.14), and a positive association with proliferation (r = 0.19), each significant in 21-23 of the 33 cohorts (Fig. 4f), whereas the corresponding associations in cell lines were positive for EMT and stemness (Section 3.2). At the single-gene level, tumor-normal changes were largely concordant across cohorts, |delta z| exceeding 0.3 in 79% of the gene-cohort pairs from the 21 cohorts with at least three adjacent normals (Fig. S1b; Fig. 4c).

Of the associations examined at the program level, the most consistent in tumor tissue was that of the composite balance score with immune programs (Fig. 4e).

### 3.4 The balance score is associated with immune infiltration and adverse outcome

An immune-cold transcriptomic state was defined here as a reduced immune program score, which is the quantity accessible from bulk expression data. Across 9,732 tumors, the balance score was inversely associated with this immune program score (pooled r = -0.24, P < 1e-100; Figs. 4e and 5f), the direction being negative in 29 of the 33 cohorts, and also with 13 individual checkpoint and effector genes: 83% of all gene-cohort pairs were negative, the strongest and most reproducible associations being found for TIGIT (mean r = -0.25; 30 of 33 cohorts), CTLA4 (r = -0.25; 31), CD8A (r = -0.23; 30) and PDCD1 (r = -0.23; 30), and the weakest for HLA-A (r = -0.04; 21) (Fig. 5e; Fig. S1e). Decomposition of the balance score into its two arms and into nine non-overlapping antioxidant sub-modules (Fig. S1g; Table S3) attributed this pattern to the generation arm rather than to antioxidant defense. ROS-generation activity correlated positively with immune infiltration in every cohort examined (per-cohort median r = +0.48 across the 32 cohorts; the pooled coefficient across all tumors is 0.49, Fig. S1c), whereas no antioxidant sub-module showed a negative association (median r between -0.02 and +0.35; the three modules closest to zero were not significant by sign test after FDR correction). Immune-cold tumors are characterized by low activity of the ROS-generation arm, and the negative balance-immune association arises arithmetically when this strongly positive term is subtracted (Fig. S1c,g).

**Figure 5.**
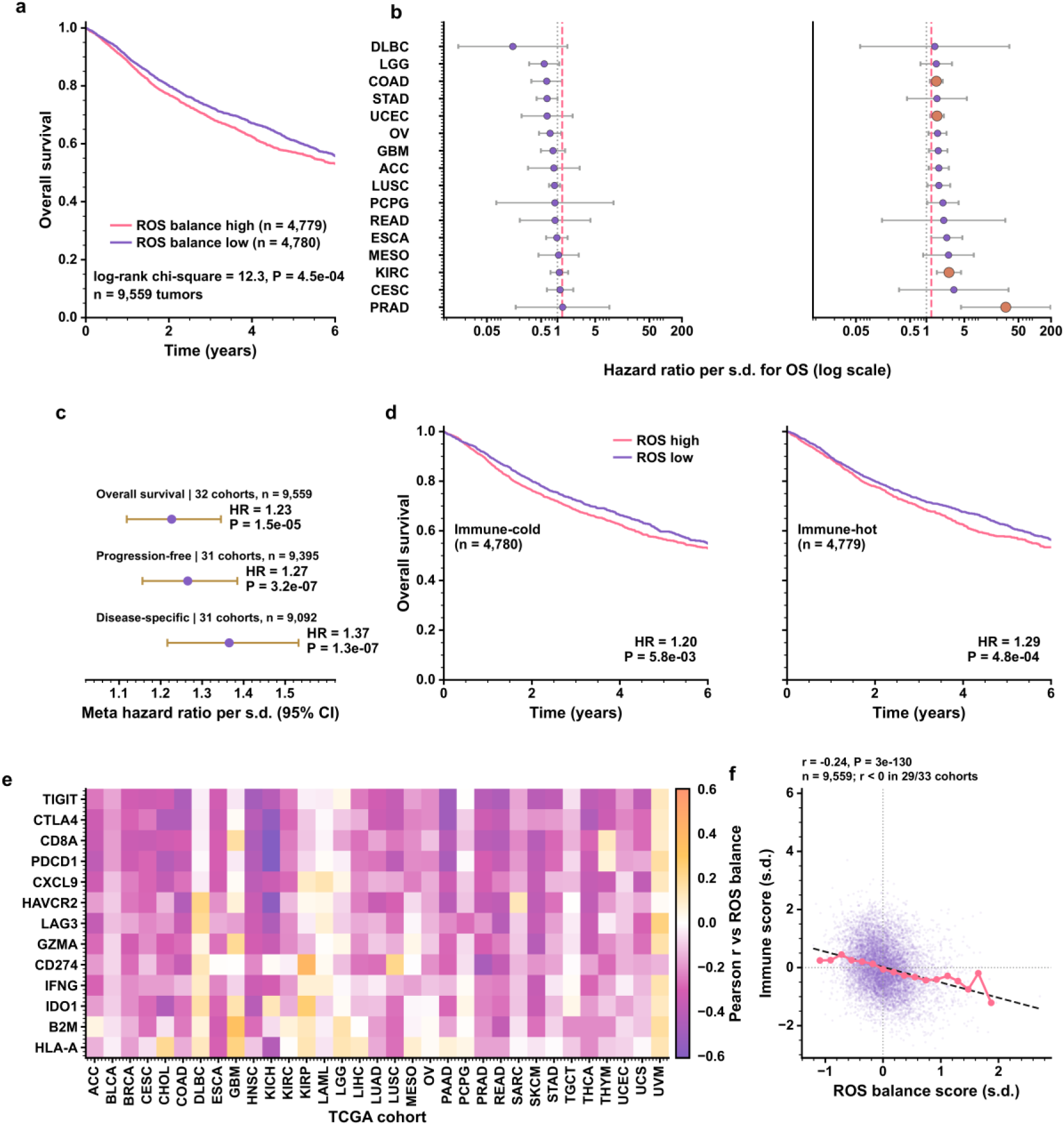
The ROS balance score, its immune correlates and adverse outcome. (a) Kaplan-Meier overall survival of 9,559 tumors stratified by the median balance score (log-rank chi-square = 12.3, P = 4.5e-04). (b) Cohort-wise univariate Cox hazard ratios per standard deviation for OS, on a log scale (32 cohorts; median HR 1.33; filled markers denote FDR < 0.05; dashed line, pooled estimate). (c) Fixed-effect meta-analysis of three survival endpoints (OS 1.23, PFI 1.27, DSS 1.37). (d) Overall survival within immune-cold and immune-hot strata defined by the median immune program score (HR 1.20 and 1.29, respectively). (e) Per-cohort correlation of the balance score with 13 immune-checkpoint and effector genes; numbers in parentheses indicate the cohorts with negative correlation. (f) The balance score versus immune program score across 9,559 tumors (r = -0.245, P < 1e-100; binned medians and linear fit shown). All panels use tumor samples only (n = 9,559 tumors, 33 cohorts).

Adverse outcome was also associated with a higher balance score. Among 9,559 tumors with survival annotation, overall survival was worse in the upper half of the balance distribution (log-rank chi-square = 12.3, P = 4.5e-04; Fig. 5a). Cohort-wise Cox models were individually underpowered but directionally consistent (median HR 1.33; HR > 1 in 62% of 32 cohorts; FDR < 0.05 in 4 cohorts; Fig. 5b), and fixed-effect meta-analysis across cohorts was significant for all three endpoints: OS HR 1.23 (95% CI 1.12-1.35, P = 1.5e-05), PFI HR 1.27 (1.16-1.39, P = 3.2e-07) and DSS HR 1.37 (1.22-1.53, P = 1.3e-07) (Fig. 5c). The association was reproduced across dichotomizations (median split, tertiles, continuous; nine models, all P < 1e-03; Fig. S1d), and the association was present in both immune-hot (HR 1.29, P = 4.8e-04) and immune-cold tumors (HR 1.20, P = 5.8e-03; Fig. 5d). Arm-level decomposition attributed this survival association to the antioxidant arm itself (univariate HR per s.d.: 1.09 for OS, P = 1.2e-06; 1.07 for PFI, P = 2.4e-04; 1.10 for DSS, P = 4.3e-05) rather than to ROS generation (HR 0.98-1.00; all P > 0.25; Fig. S1h; Table S4), and the composite balance score gave no improvement over the antioxidant score alone. The absolute hazard ratios for the composite and for the antioxidant arm differ by scale rather than by effect. The arm-level estimates (1.07-1.10 per s.d.) come from a single pan-cancer pooled model on pan-cancer standardized scores, whereas the meta-analytic estimates (1.23-1.37) combine cohort-wise models fitted on within-cohort standardized scores, in which a one-unit change corresponds to a smaller biological increment. Both decompositions place the prognostic signal in the antioxidant arm (Fig. 5c; Fig. S1h; Table S4).

The two arms of ROS homeostasis were associated with distinct aspects of tumor biology in this pan-cancer analysis. ROS-generation activity was associated with immune context, antioxidant program activity with survival, and the negative immune correlation carried by the composite balance score followed arithmetically from the low activity of the generation arm in immune-cold tumors. The two arms carry different information, and neither association arises from the antioxidant arm acting on immune infiltration.

### 3.5 Cross-cancer transferability of the ROS program

A stricter question than association is whether the same program carries information that generalizes to tumors never used in its construction. Supervised classification was applied as a test of transferability, and all validation was performed with entire cohorts held out (Fig. 6). In the 24 cohorts containing both tumors and adjacent normals (8,941 samples), cross-validated AUCs of 0.98-0.99 were obtained for all five classifiers (median cross-validated AUC: SVM 0.990, logistic regression 0.992, LASSO 0.992, gradient boosting 0.987, random forest 0.983; Fig. 6b). Under leave-one-cohort-out validation, a pooled AUC of 0.979 (95% CI 0.973-0.984) was obtained, the median cohort AUC was 0.992, and 23 of 24 cohorts exceeded 0.8 (Fig. 6c). Because per-gene standardization is performed within each cohort, including the held-out one, these predictions are cohort-held-out rather than strictly held-out quantities (Sections 2.3 and 2.8). Calibration was close to ideal (Brier score 0.023; reliability 0.0003; resolution 0.043; Fig. 6d).

**Figure 6.**
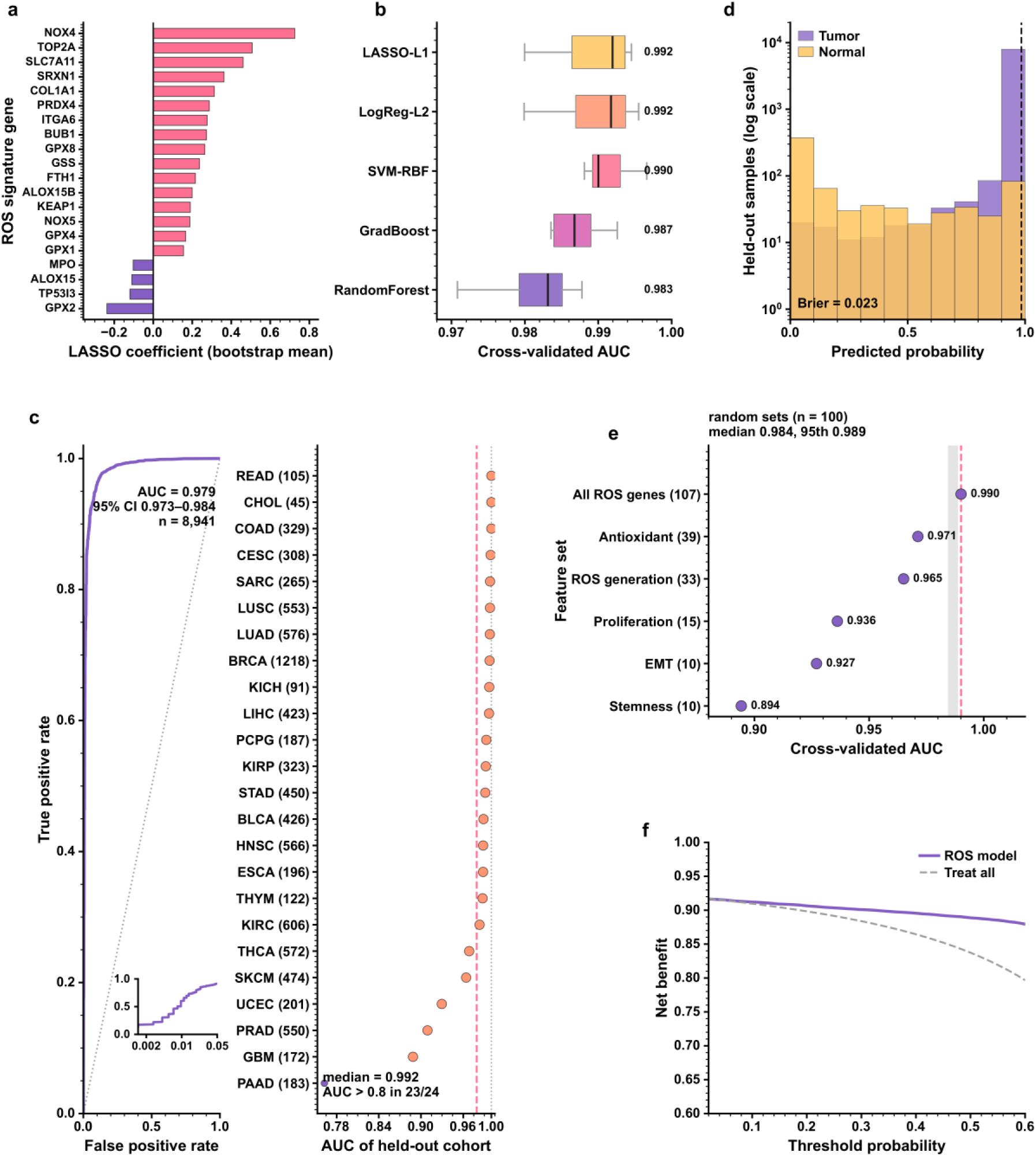
The ROS program carries information that generalizes across cancer types. (a) Signature composition from 100 bootstrap LASSO fits: LASSO coefficients (bootstrap mean) of the 20 most frequently selected genes, colored by sign (pink, positive; purple, negative); 47 of 107 genes were selected in more than half of the fits and 33 in all 100 fits. (b) Cross-validated AUC of five classifiers (stratified 5-fold cross-validation repeated three times). (c) Leave-one-cohort-out validation: pooled ROC of held-out predictions (AUC 0.979, 95% CI 0.973-0.984; 8,941 samples) and per-cohort AUCs (orange markers, AUC > 0.8 in 23 of 24 cohorts; the dashed line marks the pooled AUC). The single exception is PAAD, whose AUC rests on four adjacent normal samples. The inset re-plots the pooled ROC against a logarithmic false-positive-rate axis (0.0014-0.05; the lower limit is the smallest resolvable rate, 1/727). (d) Distribution of the held-out predicted probabilities by true class (tumor, normal; log scale), with the Brier score (0.023); the dashed line marks the Youden threshold (0.98). (e) Ablation analysis: full ROS signature (dashed line), individual program subsets (AUCs annotated in the panel) and 100 size-matched random gene sets (gray band, median to 95th percentile; none reached the full signature, empirical P = 0.010). (f) Decision curve analysis of the ROS classifier, plotted over threshold probabilities of 0.02-0.60 (the ROS classifier exceeds treat-all from 0.04 onwards): net benefit versus the treat-all reference strategy (the treat-none reference is identically zero and lies below the plotted net-benefit axis, which begins at 0.6). All panels use the same 107 ROS program genes as Figures 2-5; panels c, d and f report held-out predictions under leave-one-cohort-out training (the model was fitted on 23 cohorts and evaluated on the held-out cohort), whereas the feature-selection and classifier comparisons in panels a, b and e used bootstrap resampling and repeated stratified 5-fold cross-validation. Per-gene standardization is performed within each cohort, so panels c, d and f report cohort-held-out rather than strictly held-out predictions (Methods).

Feature selection was stable: 47 of 107 genes were selected in more than half of 100 bootstrap LASSO fits, headed by NOX4, TOP2A, SLC7A11, SRXN1, COL1A1 and PRDX4 (Fig. 6a). In ablation analysis, individual programs were sufficient but not equivalent - antioxidant genes (AUC 0.971) and generation genes (0.965) exceeded proliferation (0.936), EMT (0.927) and stemness (0.894) subsets, whereas the full signature reached 0.990 (Fig. 6e) - indicating that tumor-state information is concentrated preferentially in redox programs, and in the antioxidant arm in particular. For context, 100 size-matched random gene sets were evaluated: their median AUC was 0.984, and none reached the ROS signature (empirical P = 0.010; Fig. 6e). Although tumor-versus-normal discrimination is intrinsically an easy task, the ROS program still performed significantly better than matched random features. Net benefit above treat-all was indicated by decision curve analysis over threshold probabilities of 0.02-0.60, exceeding treat-all from 0.04 onwards (Fig. 6f), with a sensitivity of 0.990 and a specificity of 0.759 at the maximum-accuracy threshold (accuracy 0.971; Fig. S1f).

## 4. Discussion

The previously reported optical ROS fingerprint [15,16] was found to reflect a general property of human tumors. Remodeling of ROS homeostasis was coordinated across the generation and antioxidant arms, and tumor-normal differences involved the balance between the arms rather than ROS abundance alone. The most consistent association in tumor tissue was that of the balance score with reduced immune infiltration; arm-level decomposition showed that low generation-arm activity in immune-cold tumors drove this pattern and that it follows arithmetically from the definition of the balance score, a distinction that a composite score alone could not have made. Associations of the balance score with EMT, stemness and proliferation were context dependent, with opposite directions in cell lines and in tumor tissue, and are reported as accompanying phenotypic correlates. The clinical association was carried by the antioxidant arm (meta hazard ratio 1.23-1.37 across overall, progression-free and disease-specific survival) whereas the generation arm was null. The same 107-gene program transferred across cancer types, separating tumor from adjacent normal tissue in cohorts never used for training.

These observations are consistent with the concept that tumor cells convert ROS into a signaling and survival advantage through expansion of glutathione and thioredoxin metabolism [6-10]. A strengthened antioxidant arm is compatible with SLC7A11-dependent cystine uptake, limitation of lipid peroxidation and ferroptosis [10,11], and with the immune evasion and T cell exclusion that characterize such tumors [12-14]. In the present pan-cancer analysis, immune infiltration was associated with the generation arm and survival with the antioxidant arm; these associations are complementary, and redox-directed therapies would be expected to act on both arms. The survival effect was modest in magnitude but highly consistent across cohorts, endpoints and stratification schemes. Predictive value for immune-checkpoint blockade is not established: in the public immunotherapy cohorts available the direction of association was not reproducible, and the question remains open.

Program scores are expression-based proxies of the ROS state: the public cohorts contain no direct measurement of ROS or of antioxidant metabolites, and bulk tissue averages heterogeneous cell populations. Immune cells contribute little to the generation program, so immune-infiltrated samples contain a lower fraction of tumor cells and will show a lower apparent generation score. This compositional effect can account for a substantial part, possibly all, of the positive generation-immune correlation; purified or single-cell data, or a purity-adjusted model, will be required to separate the two. The array data and the transcriptomic cohorts are also not paired on the same samples, so the link between the optical fingerprint and the gene program is conceptual rather than directly calibrated.

Tumor-versus-normal discrimination is an intrinsically easy task, and the classifier z-scores genes within the cohort being scored, so those scores are not strictly held-out quantities: size-matched random gene sets already reach a median AUC close to that of the full signature, and roughly three-quarters of the program overlaps the proliferation, EMT and stemness axes. The classification results are evidence of cross-cancer transferability rather than a clinical diagnostic, and the program is not specific to redox biology. The composite balance score, being a difference of two correlated terms, likewise carries no independent coefficient interpretation, so the arm-level decomposition (Table S4) is the appropriate summary; the larger apparent hazard ratio of the composite relative to the per-standard-deviation antioxidant coefficient reflects cohort-versus pan-cancer-standardized scales and between-cohort heterogeneity.

All analyses are cross-sectional and observational, and clinical annotation varies in completeness. The phenotypic correlates of the balance score are context dependent, with opposite directions for EMT and stemness in cell lines and in tumor tissue. Causality and therapeutic direction remain to be tested.

Future work should pair optical ROS readouts with transcriptomic profiling in the same specimens, test whether antioxidant-shifted tumors respond differently to redox-directed or immune-directed therapies in prospective designs, and resolve the cell types in which the antioxidant program is induced within the tumor microenvironment.

## 5. Conclusions

Remodeling of ROS homeostasis, rather than simple elevation of ROS abundance, is a widespread feature of human tumors. This remodeling is coordinated across the generation and antioxidant arms and is measurable both optically in cells and transcriptomically across cancer types. A survival association was attributable to the antioxidant arm, and no prognostic benefit was obtained from combining the two arms. That the program transfers across cancer types places tumor remodeling of ROS homeostasis among the quantifiable features of cancer biology.

## Declarations

### Ethics approval and consent to participate

The 12 clinical specimens analyzed in this study (Section 2.1 and Fig. 2g) are the 12 cases of invasive breast cancer samples first reported in [15]. As described there, the samples were collected from the First Affiliated Hospital of Anhui Medical University, the patients’ ages ranged from 40 to 65 years, and the original study was conducted according to the regulations of the Biomedical Ethics Committee of Anhui Medical University (No. 83220390). The specimens were re-analyzed here in de-identified form; no new human material was collected for the present study.

### Availability of data and materials

This study re-analyzed previously published data an d publicly available datasets; no new experimental data were generated. The datasets use d were: (i) the AuNCs@His ROS-response array measurements first reported in [15] (11 cell lines and 12 clinical specimens; Figs. 1 and 2), with the underlying emission spectra and the archived project files available from the corresponding authors; (ii) TCGA pan-cancer expression and clinical data, obtained as the Xena-normalized HiSeqV2 log2(normal ized count + 1) matrices and the Survival_SupplementalTable_S1_20171025_xena_sp su rvival table for all 33 TCGA cohorts (https://xenabrowser.net/datapages/, Pan-Cancer Atl as hub), with sample annotation from TCGA_phenotype_denseDataOnlyDownload.tsv.g z; (iii) CCLE cell-line RNA-seq expression (RSEM TPM) from the DepMap portal (https://depmap.org/portal/, file CCLE_RNAseq_rsem_genes_tpm_20180929.txt.gz); and (iv) t he public immunotherapy cohorts GSE78220, GSE91061, GSE135222 and GSE176307 f rom NCBI GEO. For the public datasets, versions, download URLs and file checksums ar e recorded in data/public/DOWNLOAD_MANIFEST.csv. All derived data tables support ing the figures are provided as supplementary tables (Tables S1-S6).

### Code availability

All analysis and figure-generation code, together with the intermediate data tables, is available from the corresponding authors on reasonable request.

### Authors’ contributions

Kun Xie and Yulong Xuan contributed equally as co-first authors. They conceived and designed the study, defined the analysis framework and the overall implementation strategy, performed the statistical analyses and drafted the manuscript. Ruyin Chang, Jiajing Lin, Youqi Shen and Wenyue Jin contributed to data curation, generation of the figures and assembly of the supplementary tables. Rongjun Sun and Qi Lu performed the nanocluster-array experiments and the optical measurements, and contributed to data curation and interpretation.

