## Supplementary Material for "Pan-cancer remodeling of reactive oxygen species homeostasis: distinct roles of the ROS-generation and antioxidant arms in immune context and outcome"

### Supplementary information

This document contains the supplementary figure and tables for the manuscript. Panel and table captions correspond verbatim to those cited in the main text.

### Supplementary figure

##### Figure S1

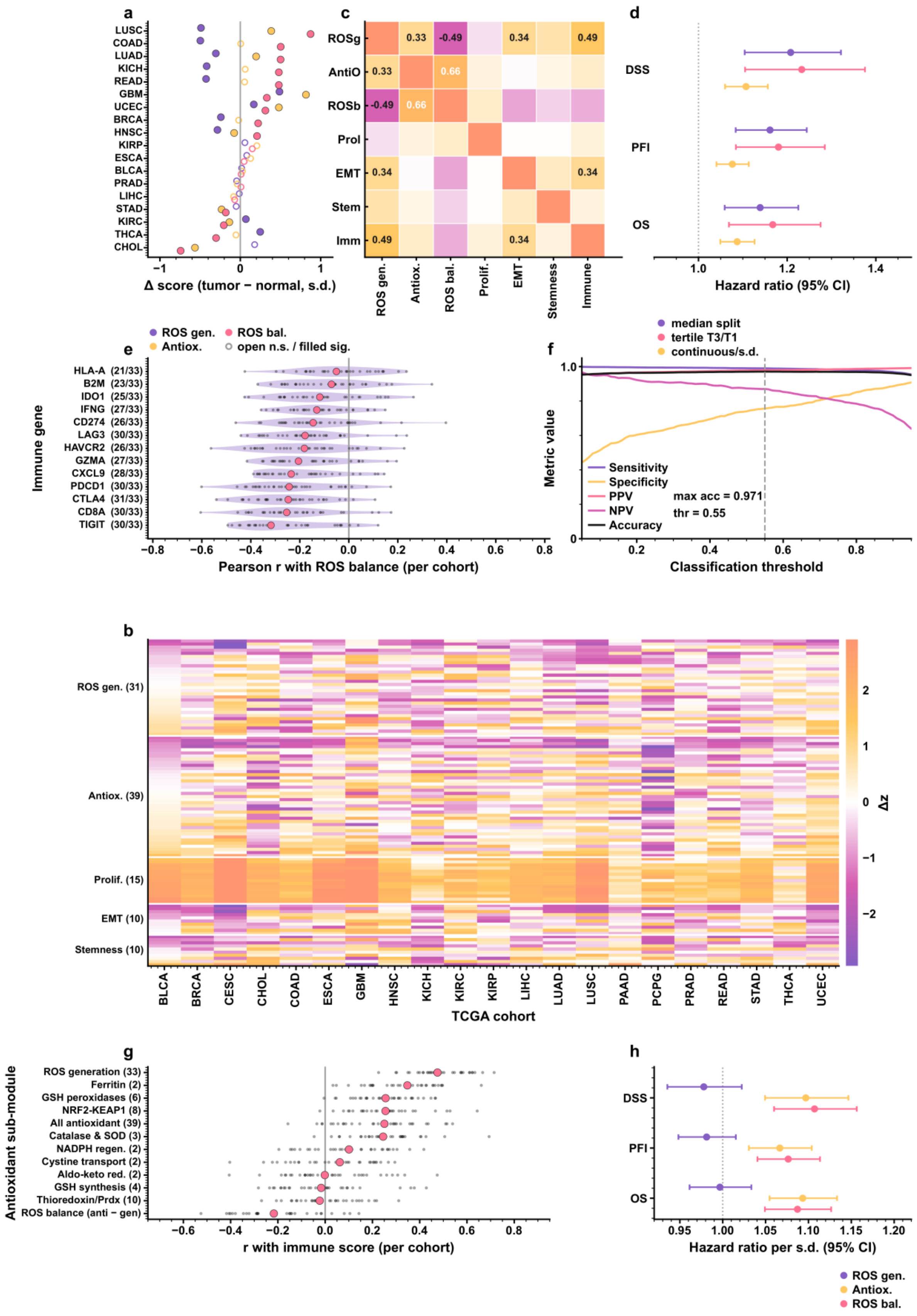


Figure S1 | Extension and robustness analyses supporting Figures 4-6. (a) Tumor minus adjacent-normal differences for the ROS generation, antioxidant and balance scores in the 18 cohorts with evaluable adjacent normal tissue (filled markers, FDR < 0.05; 11, 8 and 13 cohorts, respectively). (b) Heat map of per-gene tumor-normal differences (delta z) for 105 ROS program genes (the two remaining program entries, NOX2 and PIG3, are alternative symbols for CYBB and TP53I3, i.e. the same two genes; the program therefore comprises 105 unique genes, and the alias entries contribute to no score) in the 21 cohorts with at least three adjacent normals; 79% of gene-cohort pairs exceed |delta z| > 0.3 (Fig. 4c shows a smaller, high-variability subset of these genes). (c) Correlation matrix among the ROS, proliferation, EMT, stemness and immune program scores across all 9,732 tumors (pooled); note that ROS generation correlates positively with immune infiltration (r = 0.49), whereas the antioxidant arm dominates the negative association of ROS balance with immune score. (d) Sensitivity of the survival association to dichotomization: hazard ratios (95% CI) for median split, tertile comparison and continuous models across OS, PFI and DSS (all P < 1e-03). (e) Distribution of per-cohort correlations between ROS balance and each of 13 immune genes across 33 cohorts, with median and the number of negative cohorts annotated. (f) Operating characteristics of the ROS classifier as a function of threshold, including sensitivity, specificity, positive and negative predictive value and accuracy; the maximum-accuracy threshold is 0.55 (accuracy 0.971, sensitivity 0.990, specificity 0.759). (g) Decomposition of the antioxidant arm into nine non-overlapping sub-modules (GSH synthesis, GSH peroxidases, catalase/SOD, thioredoxin-peroxiredoxin, NRF2-KEAP1 axis, cystine transport, NADPH regeneration, ferritin, aldo-keto reductases; Table S3) plus the composite all-antioxidant score (39 genes; the parent module of the nine sub-modules), the generation arm and the composite balance score (12 rows): distribution of per-cohort correlations with the immune program score across 32 cohorts (dots, individual cohorts; bar, median). ROS generation is uniformly positive (median r = +0.48; 0/32 negative), no antioxidant sub-module is negatively associated with immune infiltration, and the negative value of the balance score is inherited arithmetically from the subtracted generation term. (h) Univariate hazard ratios per standard deviation for the two arms and the composite balance score across OS, PFI and DSS (n = 9,559 tumors; Table S4): the prognostic association is carried by the antioxidant arm, whereas ROS generation is null. Panel b uses the 21 cohorts with at least three adjacent normal samples; all statistics follow Figures 2-6.

### Supplementary tables

## S1

Table S1 | Tumor-normal differences and FDR values for the ROS generation, antioxidant and balance scores in the 18 cohorts with evaluable adjacent normal tissue.

| Cohort | Tumors (n) | Normals (n) | ROS generation delta | ROS generation FDR | Antioxidant delta | Antioxidant FDR | ROS balance delta | ROS balance FDR |
|---|---|---|---|---|---|---|---|---|
| ACC | 79 | 0 | n.a. | n.a. | n.a. | n.a. | n.a. | n.a. |
| BLCA | 407 | 19 | 0.016 | 0.765 | 0.026 | 0.600 | 0.010 | 0.891 |
| BRCA | 1104 | 114 | -0.242 | < 0.001 | -0.020 | 0.600 | 0.222 | < 0.001 |
| CESC | 305 | 3 | n.a. | n.a. | n.a. | n.a. | n.a. | n.a. |
| CHOL | 36 | 9 | 0.178 | 0.101 | -0.564 | < 0.001 | -0.743 | < 0.001 |
| COAD | 288 | 41 | -0.497 | < 0.001 | 0.005 | 0.973 | 0.503 | < 0.001 |
| DLBC | 48 | 0 | n.a. | n.a. | n.a. | n.a. | n.a. | n.a. |
| ESCA | 185 | 11 | 0.080 | 0.337 | 0.128 | 0.600 | 0.047 | 0.891 |
| GBM | 167 | 5 | 0.487 | 0.001 | 0.817 | < 0.001 | 0.330 | 0.026 |
| HNSC | 522 | 44 | -0.285 | < 0.001 | -0.077 | 0.021 | 0.208 | 0.004 |
| KICH | 66 | 25 | -0.419 | < 0.001 | 0.060 | 0.450 | 0.479 | < 0.001 |
| KIRC | 534 | 72 | 0.069 | 0.020 | -0.137 | < 0.001 | -0.206 | < 0.001 |
| KIRP | 291 | 32 | 0.057 | 0.130 | 0.206 | 0.074 | 0.149 | 0.213 |
| LAML | 173 | 0 | n.a. | n.a. | n.a. | n.a. | n.a. | n.a. |
| LGG | 530 | 0 | n.a. | n.a. | n.a. | n.a. | n.a. | n.a. |
| LIHC | 373 | 50 | -0.013 | 0.734 | -0.083 | 0.101 | -0.070 | 0.318 |
| LUAD | 517 | 59 | -0.305 | < 0.001 | 0.196 | 0.001 | 0.501 | < 0.001 |
| LUSC | 502 | 51 | -0.490 | < 0.001 | 0.385 | < 0.001 | 0.874 | < 0.001 |
| MESO | 87 | 0 | n.a. | n.a. | n.a. | n.a. | n.a. | n.a. |
| OV | 308 | 0 | n.a. | n.a. | n.a. | n.a. | n.a. | n.a. |
| PAAD | 179 | 4 | n.a. | n.a. | n.a. | n.a. | n.a. | n.a. |
| PCPG | 184 | 3 | n.a. | n.a. | n.a. | n.a. | n.a. | n.a. |
| PRAD | 498 | 52 | -0.046 | 0.378 | -0.038 | 0.464 | 0.008 | 0.891 |
| READ | 95 | 10 | -0.426 | < 0.001 | 0.053 | 0.600 | 0.479 | < 0.001 |

| | | | | | | | | |
|---|---|---|---|---|---|---|---|---|
| SARC | 263 | 2 | n.a. | n.a. | n.a. | n.a. | n.a. | n.a. |
| SKCM | 473 | 1 | n.a. | n.a. | n.a. | n.a. | n.a. | n.a. |
| STAD | 415 | 35 | -0.052 | 0.130 | -0.233 | 0.001 | -0.181 | 0.009 |
| TGCT | 156 | 0 | n.a. | n.a. | n.a. | n.a. | n.a. | n.a. |
| THCA | 513 | 59 | 0.247 | < 0.001 | -0.054 | 0.130 | -0.302 | < 0.001 |
| THYM | 120 | 2 | n.a. | n.a. | n.a. | n.a. | n.a. | n.a. |
| UCEC | 177 | 24 | 0.168 | 0.005 | 0.480 | < 0.001 | 0.312 | < 0.001 |
| UCS | 57 | 0 | n.a. | n.a. | n.a. | n.a. | n.a. | n.a. |
| UVM | 80 | 0 | n.a. | n.a. | n.a. | n.a. | n.a. | n.a. |

## S2

Table S2 | The 107 ROS program genes and their program membership (33 ROS generation, 39 antioxidant, 15 proliferation, 10 EMT, 10 stemness). Two of the 107 entries are alternative symbols (NOX2 = CYBB, PIG3 = TP53I3), giving 105 unique genes.

| **Program** | **Gene** |
|---|---|
| Antioxidant | AKR1B10 |
| Antioxidant | AKR1C1 |
| Antioxidant | BACH1 |
| Antioxidant | CAT |
| Antioxidant | FTH1 |
| Antioxidant | FTL |
| Antioxidant | G6PD |
| Antioxidant | GCLC |
| Antioxidant | GCLM |
| Antioxidant | GPX1 |
| Antioxidant | GPX2 |
| Antioxidant | GPX3 |
| Antioxidant | GPX4 |
| Antioxidant | GPX7 |
| Antioxidant | GPX8 |
| Antioxidant | GSR |
| Antioxidant | GSS |
| Antioxidant | HMOX1 |
| Antioxidant | HMOX2 |
| Antioxidant | KEAP1 |
| Antioxidant | NFE2L2 |
| Antioxidant | NQO1 |
| Antioxidant | PGD |

| | |
|---|---|
| Antioxidant | PRDX1 |
| Antioxidant | PRDX2 |
| Antioxidant | PRDX3 |
| Antioxidant | PRDX4 |
| Antioxidant | PRDX5 |
| Antioxidant | PRDX6 |
| Antioxidant | SLC3A2 |
| Antioxidant | SLC7A11 |
| Antioxidant | SOD1 |
| Antioxidant | SOD2 |
| Antioxidant | SRXN1 |
| Antioxidant | TXN |
| Antioxidant | TXN2 |
| Antioxidant | TXNIP |
| Antioxidant | TXNRD1 |
| Antioxidant | TXNRD2 |
| EMT | CDH2 |
| EMT | COL1A1 |
| EMT | FN1 |
| EMT | SNAI1 |
| EMT | SNAI2 |
| EMT | TGFB1 |
| EMT | TWIST1 |
| EMT | VIM |
| EMT | ZEB1 |
| EMT | ZEB2 |
| Proliferation | AURKA |

| | |
|---|---|
| Proliferation | AURKB |
| Proliferation | BIRC5 |
| Proliferation | BUB1 |
| Proliferation | CCNA2 |
| Proliferation | CCNB1 |
| Proliferation | CDC20 |
| Proliferation | CDK1 |
| Proliferation | MKI67 |
| Proliferation | PCNA |
| Proliferation | PLK1 |
| Proliferation | RRM2 |
| Proliferation | TOP2A |
| Proliferation | TYMS |
| Proliferation | UBE2C |
| ROS generation | ACOX1 |
| ROS generation | ALOX12 |
| ROS generation | ALOX15 |
| ROS generation | ALOX15B |
| ROS generation | ALOX5 |
| ROS generation | COX4I1 |
| ROS generation | COX5A |
| ROS generation | CYBA |
| ROS generation | CYBB |
| ROS generation | DUOX1 |
| ROS generation | DUOX2 |
| ROS generation | EPX |
| ROS generation | LOX |

| | |
|---|---|
| ROS generation | MAOA |
| ROS generation | MAOB |
| ROS generation | MPO |
| ROS generation | NCF1 |
| ROS generation | NCF2 |
| ROS generation | NCF4 |
| ROS generation | NOS2 |
| ROS generation | NOS3 |
| ROS generation | NOX1 |
| ROS generation | NOX2 |
| ROS generation | NOX3 |
| ROS generation | NOX4 |
| ROS generation | NOX5 |
| ROS generation | NOXA1 |
| ROS generation | NOXO1 |
| ROS generation | PIG3 |
| ROS generation | SDHB |
| ROS generation | SOD3 |
| ROS generation | TP53I3 |
| ROS generation | XDH |
| Stemness | ALDH1A1 |
| Stemness | EPCAM |
| Stemness | ITGA6 |
| Stemness | KLF4 |
| Stemness | MYC |
| Stemness | NANOG |
| Stemness | POU5F1 |

| | |
|---|---|
| Stemness | PROM1 |
| Stemness | SOX2 |
| Stemness | THY1 |

## S3

Table S3 | The nine antioxidant sub-modules and their per-cohort correlations with the immune program score (32 cohorts; median, direction counts, sign-test P and FDR), together with the generation arm and the composite balance score as references.

| Module | Genes (n) | Cohorts (n) | Median r (vs immune) | Mean r | Cohorts with r < 0 | Fraction with r < 0 | Median r (vs 13-gene immune composite) | Sign test P | Sign test FDR |
|---|---|---|---|---|---|---|---|---|---|
| ROS balance (difference) | 0 | 32 | -0.217 | -0.205 | 28 | 0.875 | -0.239 | < 0.001 | < 0.001 |
| Thioredoxin/peroxiredoxin | 10 | 32 | -0.023 | 0.004 | 17 | 0.531 | -0.012 | 0.860 | 0.938 |
| GSH synthesis & recy-cling | 4 | 32 | -0.017 | -0.008 | 18 | 0.562 | 0.022 | 0.597 | 0.716 |
| Aldo-keto reductases | 2 | 32 | -0.002 | 0.023 | 16 | 0.500 | 0.006 | 1 | 1 |
| Cystine transport (SLC7A11) | 2 | 32 | 0.062 | 0.029 | 13 | 0.406 | 0.074 | 0.377 | 0.503 |
| NADPH regeneration | 2 | 32 | 0.101 | 0.102 | 9 | 0.281 | 0.092 | 0.020 | 0.030 |
| Catalase & SOD | 3 | 32 | 0.245 | 0.256 | 1 | 0.031 | 0.266 | < 0.001 | < 0.001 |
| All antioxidant (39) | 39 | 32 | 0.251 | 0.221 | 4 | 0.125 | 0.258 | < 0.001 | < 0.001 |
| NRF2-KEAP1 axis | 8 | 32 | 0.255 | 0.258 | 1 | 0.031 | 0.289 | < 0.001 | < 0.001 |
| GSH peroxidases (GPX) | 6 | 32 | 0.256 | 0.253 | 1 | 0.031 | 0.236 | < 0.001 | < 0.001 |
| Iron storage (ferritin) | 2 | 32 | 0.348 | 0.326 | 0 | 0 | 0.370 | < 0.001 | < 0.001 |
| ROS generation (33) | 33 | 32 | 0.475 | 0.489 | 0 | 0 | 0.500 | < 0.001 | < 0.001 |

## S4

Table S4 | Arm-level univariate Cox hazard ratios per standard deviation for ROS generation, antioxidant and balance scores across OS, PFI and DSS (n = 9,559 tumors).

| Endpoint | ROS score | Tumors (n) | HR per s.d. | 95% CI low | 95% CI high | P |
|---|---|---|---|---|---|---|
| OS | ROS generation | 9559 | 0.997 | 0.961 | 1.033 | 0.858 |
| OS | Antioxidant | 9559 | 1.093 | 1.055 | 1.133 | < 0.001 |
| OS | ROS balance | 9559 | 1.087 | 1.049 | 1.127 | < 0.001 |
| PFI | ROS generation | 9395 | 0.981 | 0.948 | 1.015 | 0.276 |
| PFI | Antioxidant | 9395 | 1.067 | 1.030 | 1.104 | < 0.001 |
| PFI | ROS balance | 9395 | 1.076 | 1.040 | 1.113 | < 0.001 |
| DSS | ROS generation | 9092 | 0.978 | 0.935 | 1.022 | 0.323 |
| DSS | Antioxidant | 9092 | 1.097 | 1.049 | 1.147 | < 0.001 |
| DSS | ROS balance | 9092 | 1.107 | 1.060 | 1.156 | < 0.001 |

**S5**

Table S5 | Complete gene-set membership for all three scoring systems: the 107-gene ROS program, the 20-gene interferon/cytotoxic immune program and the 13 checkpoint and effector genes.

| Set | Program | Gene |
|---|---|---|
| ROS program (107 genes) | ROS generation | NOX1 |
| ROS program (107 genes) | ROS generation | NOX2 |
| ROS program (107 genes) | ROS generation | CYBB |
| ROS program (107 genes) | ROS generation | NOX3 |
| ROS program (107 genes) | ROS generation | NOX4 |
| ROS program (107 genes) | ROS generation | NOX5 |
| ROS program (107 genes) | ROS generation | DUOX1 |
| ROS program (107 genes) | ROS generation | DUOX2 |
| ROS program (107 genes) | ROS generation | NOXA1 |
| ROS program (107 genes) | ROS generation | NOXO1 |
| ROS program (107 genes) | ROS generation | CYBA |
| ROS program (107 genes) | ROS generation | NCF1 |
| ROS program (107 genes) | ROS generation | NCF2 |
| ROS program (107 genes) | ROS generation | NCF4 |
| ROS program (107 genes) | ROS generation | XDH |
| ROS program (107 genes) | ROS generation | MPO |
| ROS program (107 genes) | ROS generation | EPX |
| ROS program (107 genes) | ROS generation | ALOX5 |
| ROS program (107 genes) | ROS generation | ALOX12 |
| ROS program (107 genes) | ROS generation | ALOX15 |
| ROS program (107 genes) | ROS generation | ALOX15B |
| ROS program (107 genes) | ROS generation | MAOA |
| ROS program (107 genes) | ROS generation | MAOB |

| | | |
|---|---|---|
| ROS program (107 genes) | ROS generation | NOS2 |
| ROS program (107 genes) | ROS generation | NOS3 |
| ROS program (107 genes) | ROS generation | COX4I1 |
| ROS program (107 genes) | ROS generation | COX5A |
| ROS program (107 genes) | ROS generation | SDHB |
| ROS program (107 genes) | ROS generation | ACOX1 |
| ROS program (107 genes) | ROS generation | SOD3 |
| ROS program (107 genes) | ROS generation | LOX |
| ROS program (107 genes) | ROS generation | PIG3 |
| ROS program (107 genes) | ROS generation | TP53I3 |
| ROS program (107 genes) | Antioxidant | SOD1 |
| ROS program (107 genes) | Antioxidant | SOD2 |
| ROS program (107 genes) | Antioxidant | CAT |
| ROS program (107 genes) | Antioxidant | GPX1 |
| ROS program (107 genes) | Antioxidant | GPX2 |
| ROS program (107 genes) | Antioxidant | GPX3 |
| ROS program (107 genes) | Antioxidant | GPX4 |
| ROS program (107 genes) | Antioxidant | GPX7 |
| ROS program (107 genes) | Antioxidant | GPX8 |
| ROS program (107 genes) | Antioxidant | PRDX1 |
| ROS program (107 genes) | Antioxidant | PRDX2 |
| ROS program (107 genes) | Antioxidant | PRDX3 |
| ROS program (107 genes) | Antioxidant | PRDX4 |
| ROS program (107 genes) | Antioxidant | PRDX5 |
| ROS program (107 genes) | Antioxidant | PRDX6 |
| ROS program (107 genes) | Antioxidant | TXN |
| ROS program (107 genes) | Antioxidant | TXN2 |

| | | |
|---|---|---|
| ROS program (107 genes) | Antioxidant | TXNRD1 |
| ROS program (107 genes) | Antioxidant | TXNRD2 |
| ROS program (107 genes) | Antioxidant | TXNIP |
| ROS program (107 genes) | Antioxidant | GCLC |
| ROS program (107 genes) | Antioxidant | GCLM |
| ROS program (107 genes) | Antioxidant | GSR |
| ROS program (107 genes) | Antioxidant | GSS |
| ROS program (107 genes) | Antioxidant | NQO1 |
| ROS program (107 genes) | Antioxidant | HMOX1 |
| ROS program (107 genes) | Antioxidant | HMOX2 |
| ROS program (107 genes) | Antioxidant | NFE2L2 |
| ROS program (107 genes) | Antioxidant | KEAP1 |
| ROS program (107 genes) | Antioxidant | SLC7A11 |
| ROS program (107 genes) | Antioxidant | SLC3A2 |
| ROS program (107 genes) | Antioxidant | G6PD |
| ROS program (107 genes) | Antioxidant | PGD |
| ROS program (107 genes) | Antioxidant | SRXN1 |
| ROS program (107 genes) | Antioxidant | AKR1C1 |
| ROS program (107 genes) | Antioxidant | AKR1B10 |
| ROS program (107 genes) | Antioxidant | FTH1 |
| ROS program (107 genes) | Antioxidant | FTL |
| ROS program (107 genes) | Antioxidant | BACH1 |
| ROS program (107 genes) | Proliferation | MKI67 |
| ROS program (107 genes) | Proliferation | PCNA |
| ROS program (107 genes) | Proliferation | TOP2A |
| ROS program (107 genes) | Proliferation | CCNB1 |
| ROS program (107 genes) | Proliferation | CCNA2 |

| | | |
|---|---|---|
| ROS program (107 genes) | Proliferation | CDK1 |
| ROS program (107 genes) | Proliferation | CDC20 |
| ROS program (107 genes) | Proliferation | BUB1 |
| ROS program (107 genes) | Proliferation | AURKA |
| ROS program (107 genes) | Proliferation | AURKB |
| ROS program (107 genes) | Proliferation | PLK1 |
| ROS program (107 genes) | Proliferation | TYMS |
| ROS program (107 genes) | Proliferation | RRM2 |
| ROS program (107 genes) | Proliferation | UBE2C |
| ROS program (107 genes) | Proliferation | BIRC5 |
| ROS program (107 genes) | EMT | VIM |
| ROS program (107 genes) | EMT | CDH2 |
| ROS program (107 genes) | EMT | FN1 |
| ROS program (107 genes) | EMT | SNAI1 |
| ROS program (107 genes) | EMT | SNAI2 |
| ROS program (107 genes) | EMT | TWIST1 |
| ROS program (107 genes) | EMT | ZEB1 |
| ROS program (107 genes) | EMT | ZEB2 |
| ROS program (107 genes) | EMT | TGFB1 |
| ROS program (107 genes) | EMT | COL1A1 |
| ROS program (107 genes) | Stemness | PROM1 |
| ROS program (107 genes) | Stemness | SOX2 |
| ROS program (107 genes) | Stemness | NANOG |
| ROS program (107 genes) | Stemness | POU5F1 |
| ROS program (107 genes) | Stemness | ALDH1A1 |
| ROS program (107 genes) | Stemness | KLF4 |
| ROS program (107 genes) | Stemness | MYC |

| | | |
|---|---|---|
| ROS program (107 genes) | Stemness | THY1 |
| ROS program (107 genes) | Stemness | ITGA6 |
| ROS program (107 genes) | Stemness | EPCAM |
| Immune program (20 genes) | Immune | CD8A |
| Immune program (20 genes) | Immune | CD8B |
| Immune program (20 genes) | Immune | GZMA |
| Immune program (20 genes) | Immune | GZMB |
| Immune program (20 genes) | Immune | PRF1 |
| Immune program (20 genes) | Immune | IFNG |
| Immune program (20 genes) | Immune | CXCL9 |
| Immune program (20 genes) | Immune | CXCL10 |
| Immune program (20 genes) | Immune | PDCD1 |
| Immune program (20 genes) | Immune | CD274 |
| Immune program (20 genes) | Immune | CTLA4 |
| Immune program (20 genes) | Immune | LAG3 |
| Immune program (20 genes) | Immune | HAVCR2 |
| Immune program (20 genes) | Immune | TIGIT |
| Immune program (20 genes) | Immune | CD3D |
| Immune program (20 genes) | Immune | CD3E |
| Immune program (20 genes) | Immune | NKG7 |
| Immune program (20 genes) | Immune | GNLY |
| Immune program (20 genes) | Immune | STAT1 |
| Immune program (20 genes) | Immune | IRF1 |
| Immune checkpoint/effector (13 genes) | Immune checkpoint/effector | PDCD1 |
| Immune checkpoint/effector (13 genes) | Immune checkpoint/effector | CD274 |
| Immune checkpoint/effector (13 genes) | Immune checkpoint/effector | CTLA4 |
| Immune checkpoint/effector (13 genes) | Immune checkpoint/effector | LAG3 |

| | | |
|---|---|---|
| Immune checkpoint/effector (13 genes) | Immune checkpoint/effector | HAVCR2 |
| Immune checkpoint/effector (13 genes) | Immune checkpoint/effector | TIGIT |
| Immune checkpoint/effector (13 genes) | Immune checkpoint/effector | IDO1 |
| Immune checkpoint/effector (13 genes) | Immune checkpoint/effector | CD8A |
| Immune checkpoint/effector (13 genes) | Immune checkpoint/effector | IFNG |
| Immune checkpoint/effector (13 genes) | Immune checkpoint/effector | GZMA |
| Immune checkpoint/effector (13 genes) | Immune checkpoint/effector | CXCL9 |
| Immune checkpoint/effector (13 genes) | Immune checkpoint/effector | B2M |
| Immune checkpoint/effector (13 genes) | Immune checkpoint/effector | HLA-A |

## S6

Table S6 | Array classification in the 55-sample, 11 cell-line panel: full leave-one-out confusion matrix, per-model accuracy, macro-F1 and bootstrap 95% confidence intervals, and the permutation test.

##### (i) Leave-one-out confusion matrix

| True \ Predicted | A549 | HCA-EC | HK-2 | Hela | HepG2 | Het-1A | L-02 | MBA-MB-231 | MGC-803 | SKOV-3 | hOMF |
|---|---|---|---|---|---|---|---|---|---|---|---|
| A549 | 5 | 0 | 0 | 0 | 0 | 0 | 0 | 0 | 0 | 0 | 0 |
| HCA-EC | 0 | 5 | 0 | 0 | 0 | 0 | 0 | 0 | 0 | 0 | 0 |
| HK-2 | 0 | 0 | 5 | 0 | 0 | 0 | 0 | 0 | 0 | 0 | 0 |
| Hela | 0 | 0 | 0 | 5 | 0 | 0 | 0 | 0 | 0 | 0 | 0 |
| HepG2 | 0 | 0 | 0 | 0 | 5 | 0 | 0 | 0 | 0 | 0 | 0 |
| Het-1A | 0 | 0 | 0 | 0 | 0 | 5 | 0 | 0 | 0 | 0 | 0 |
| L-02 | 0 | 0 | 0 | 0 | 0 | 0 | 5 | 0 | 0 | 0 | 0 |
| MBA-MB-231 | 0 | 0 | 0 | 0 | 0 | 0 | 0 | 5 | 0 | 0 | 0 |
| MGC-803 | 0 | 0 | 0 | 0 | 0 | 0 | 0 | 0 | 5 | 0 | 0 |
| SKOV-3 | 0 | 0 | 0 | 0 | 0 | 0 | 0 | 0 | 0 | 5 | 0 |
| hOMF | 0 | 0 | 0 | 0 | 0 | 0 | 0 | 0 | 0 | 0 | 5 |

##### (ii) Per-model performance and bootstrap 95% CI

| Model | LOO accuracy | Macro F1 | Samples (n) | Classes (n) | LOO accuracy (recomputed) | 95% CI low | 95% CI high |
|---|---|---|---|---|---|---|---|
| LDA | 1.000 | 1.000 | 55 | 11 | 1.000 | 1.000 | 1.000 |
| RandomForest | 1.000 | 1.000 | 55 | 11 | 1.000 | 1.000 | 1.000 |
| SVM-RBF | 1.000 | 1.000 | 55 | 11 | 1.000 | 0.932 | 1.000 |
| kNN(k=3) | 1.000 | 1.000 | 55 | 11 | 1.000 | 0.977 | 1.000 |

##### (iii) Permutation test

| Statistic | Value |
| --- | --- |
| Permutations (n) | 200 |
| Null mean accuracy | 0.057 |
| Null 95th percentile accuracy | 0.127 |
| P | 0.005 |
